# Probing the network structure of health deficits in human aging

**DOI:** 10.1101/270876

**Authors:** Spencer G. Farrell, Arnold B. Mitnitski, Olga Theou, Kenneth Rockwood, Andrew D. Rutenberg

## Abstract

Human aging leads to the stochastic accumulation of damage. We model an aging population using a stochastic network model. Individuals are modeled as a network of interacting nodes, representing health attributes. Nodes in the network stochastically damage and repair, with rates dependent on the state of their neighbors. Damaged nodes represent health deficits. The Frailty Index (FI) assesses age-related damage as the proportion of health deficits an individual has accumulated, from a selection of attributes. Here, we use computational, information-theoretic, and mean-field approaches to show that the degree distribution and degree correlations of the underlying network are important to the model’s ability to recover the behavior of observational health data. We use different measures of damage in the network to probe the structure of the network. We find that the behavior of different classes of observational health deficits (laboratory or clinical) is similar to the behavior of nodes of low or high degree in the model, respectively. This explains how damage can propagate within the network, leading towards individual mortality.

## I. INTRODUCTION

Accumulation of damage causes organismal aging [1]. Even in model organisms, with controlled environment and genotype, there are large individual variations in lifespan and in the phenotypes of aging [2, 3]. While many mechanisms cause specific cellular damage [4], no single factor fully controls the process of aging. This suggests that the aging process is stochastic and results from a variety of damage mechanisms.

The variability of individual damage accumulation results in differing trajectories of individual health and in differing individual lifespans, and is a fundamental aspect of individual aging. A simple method of quantifying this individual damage is the Frailty Index (FI) [5, 6]. The FI is the proportion of agerelated health issues (“deficits”) that a person has out of a collection of health attributes. The FI is used as a quantitative tool in understanding the health of individuals as they age in clinical and epidemiological studies. There have been hundreds of papers using an FI based on self-report or clinical data, both for humans [7] and for animals [8]. Individuals typically accumulate deficits as they age, and so the FI increases with age across a population. The FI captures the heterogeneity in individual health and is predictive of both mortality and other health outcomes [9–11].

In previous work we developed a stochastic network model of aging with damage accumulation [12, 13]. Each individual is modeled as a network of interacting nodes that represent health attributes. Both the nodes and their connections are idealized and do not specify particular health aspects or mechanisms. Connections (links) between neighboring nodes in the network can be interpreted as influence between separate physiological systems. In our model, damage facilitates subsequent damage of connected nodes. We do not specify the biological mechanisms that cause damage, only that damage rates depend on the proportion of damaged neighbors. Damage promotes more damage and lack of damage facilitates repair. Rather than model the specific biological mechanisms of aging, we model how damage to components of generic physiological systems can accumulate and propagate throughout an organism — ending with death.

Even though our model includes no explicit age-dependence in damage rates or mortality, it captures Gompertz’s law of mortality [14, 15], the increase average rate of FI accumulation [5], 16], and the broadening of FI distributions with age [17, 18]. By including a falsenegative attribution error (i.e. a finite sensitivity) [13], we can also explain an empirical maximum of observed FI values – typically between 0.6 − 0.8 [6, 16–20]. This shows that age-dependent “programming” of either mortality or damage rates are not necessary to explain these features [1].

Recently, the FI approach has been extended to laboratory [21] and biomarker data [22] and used in clinical [23, 24] and population settings [25]. Two different FIs have been constructed to measure different types of damage, *F*_clin_, with clinically evaluated or selfreported data, and *F*_lab_, with lab or biomarker data. Clinical deficits are typically based on disabilities, loss of function, or diagnosis of disease, and they measure clinically observable damage that typically occurs late in life. Lab deficits or biomarkers use the results of lab tests (e.g. blood tests or vital signs) that are binarized using standard reference ranges [26]. Since frailty indices based on laboratory tests measure pre-clinical damage, they are expected to be distinct from those based on clinical and/or self-report data
[21, 25].

Even though they measure very different types of damage, both FIs are similarly associated with mortality [21, 27]. Earlier observational studies have found (average) 〈*F*_lab_〉 larger than 〈*F*_clin_〉 [21, 22, 27]. However, a study of older long-term care patients has found 〈*F*_lab_〉 less than 〈*F*_clin_〉 [28]. While differences between studies could be attributed to classification differences, a large single study including ages from 20-85 from the National Health and Nutrition Examination Survey (NHANES) [25] found that 〈*F*_lab_〉 was higher than 〈*F*_clin_〉 at earlier ages, but below at later ages.

The observed age-dependent relationship (or “age-structure”) between *F*_lab_ and *F*_clin_ challenges us to ex-amine whether network properties can provide similar age-structure in model data. Previously, we had chosen the Barabási-Albert (BA) preferential attachment algorithm [29] to generate our scale-free network, both due to the simplicity of the BA algorithm and due to the numerous examples of these scale-free networks in biological systems [30].

Allowing age-dependent damage rates to differ for different network nodes would significantly increase the parameter freedom of our model, and let us provide facile agreement with the observed phenomenon. Instead, we hope that the network topology itself is sufficient to recover the observed age-structure without requiring model parameters to vary within the network. Specifically, we aim to determine what qualitative network features are necessary to explain age-structure. Our working hypothesis is that low-degree nodes should correspond to *F*_lab_, just as high-degree nodes correspond to *F*_clin_ [12, 13].

The direct assessment of node connectivity from observational data is a challenging and generally unsolved problem. Nevertheless, we can reliably reconstruct the relative connectivity of high degree nodes in both model and in large-cohort observational data by measuring mutual dependence between pairs of nodes. This reconstruction allows us to qualitatively confirm the relationship between the connectivity of nodes and how informative they were about mortality [13].

Of course, complex networks have structural features beyond the degree distribution. For example, degree correlations describe how connections are made between specificc nodes of diffierent degree [31]. Accordingly, we consider networks with three types of degree correlations: assortative, disassortative, and neutral [31, 32]. Networks with assortative correlations tend to connect like-degree nodes, those with disassortative correlations tend to connect unlike-degrees, and those with neutral correlations are random. We probe and understand the internal structure of these networks by examining *F*_high_ and *F*_low_, i.e. damage to high degree nodes and damage to low degree nodes.

A fundamental unanswered question of aging is how does internal network structure affect aging? We show how network properties of degree distribution and degree correlations are essential for our model to recover results from observational data. Doing so, we can explain how damage propagates through our network and what makes nodes informative of mortality. This allows us to understand the differences between *F*_low_ and *F*_high_, or between pre-clinical and clinical damage in observational health data.

## II. MODEL AND ANALYSIS

### A. Model

Our model was previously presented [13]. Individuals are represented as a network consisting of *N* nodes, where each node *i* ∈ {1, 2,…, *N*} can take on binary values *d*_*i*_ = 0,1 for healthy or damaged, respectively. Connections are undirected and all nodes are undamaged at time *t* = 0.

A stochastic process transitions between healthy and damaged (*d*_*i*_ = 0, 1) states. Healthy nodes damage with rate Γ_+_ = Γ_0_ exp (*f*_*i*_γ+) and damaged nodes repair with rate Γ_−_ = (Γ_0_/*R*) exp (−*f*_*i*_γ_−_). These rates depend on the local frailty 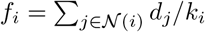, which is the proportion of damaged neighbors of node *i*. This *f*_*i*_ is like an FI but quantifies local damage within the network. Transitions between the damaged and healthy states of nodes are implemented exactly using a stochastic simulation algorithm [33, 34]. Individual mortality occurs when the two highest degree nodes are both damaged.

We generate our default network “topology” using a linearly-shifted preferential attachment algorithm [35, 36], which is a generalization of the original Barabási Albert algorithm [29]. This generates a scale-free network *P*(*k*) ~ *k*^−α^, where the exponent *α* and average degree 〈k〉 can be tuned. (The minimum degree varies as *k*_min_ = 〈*k*〉/2.) This network is highly heterogeneous in both degree *k*_*i*_ and nearest-neighbor degree (nn-degree) 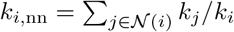.

Since we are concerned with the properties of individual nodes and groups of nodes, we use the same randomly generated network for all individuals. As a result, connections between any two nodes are the same for every individual. To ensure that our randomly generated network is generic, we then redo all of our analysis for 10 different randomly generated networks. We find insignificant differences between the networks, and so present results averaged over them. Previously [13], we had randomly generated a distinct network realization for each individual.

We use observational data for mortality rate and FI vs age to fine-tune the network parameters [12, 13]. A systematic exploration of parameters was done in previous work [12, 13]. Most of our parameterization (*N* = 10000, α = 2.27, 〈*k*〉 = 4, γ_−_ = 6.5) is the same as reported previously [13]. However, three parameters (Γ_0_ = 0.00183/yr, γ_+_ = 7.5, *R* = 3) have been adjusted because we now disallow multiple connections between nodes in our network generation. This eases analysis and adjustment of the network topology but would affect mortality rates (see e.g. Fig. 15 in Appendix A below) without the parameter adjustment. Other network topologies, explored and described below in Sect. III D, use this same “default” parameterization.

**FIG. 1.**
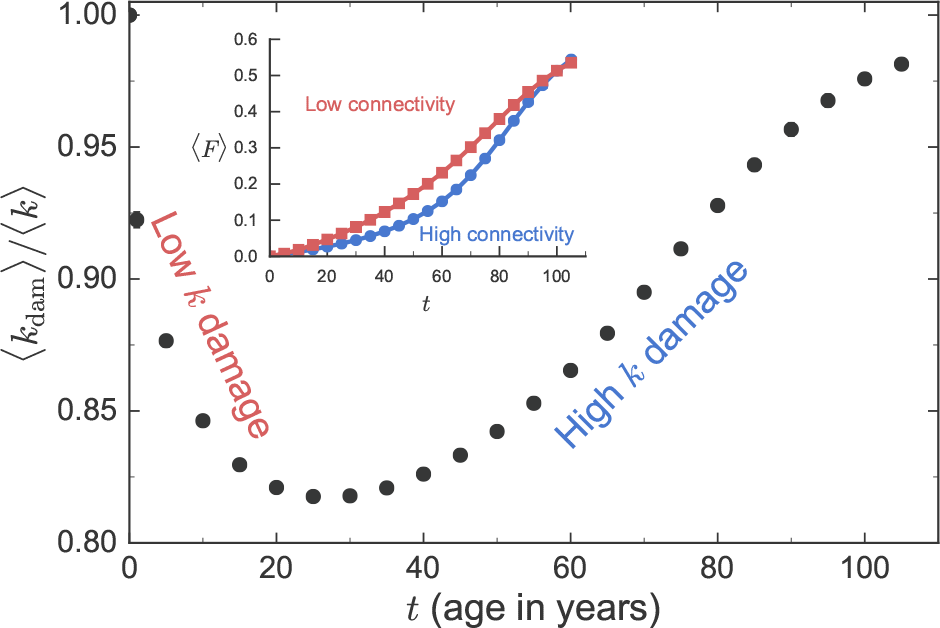
Average degree of damaged model deficits 〈*k*_dam_(*t*)〉, scaled by the average degree of the network 〈k〉, vs time *t*. Error bars, barely visible at low *t*, represent the standard deviation between randomly generated networks. As indicated, at earlier times low-connectivity nodes are preferentially damaged while at later times higher connectivity nodes are preferentially damaged. The inset shows the average damage of low-connectivity nodes 〈*F*_low_〉 (red squares) and of high-connectivity nodes 〈*F*_high_〉 (blue circles) vs age.

### B. Analysis

Typically, binary deficits have a finite sensitivity [37] while our model gives us exact knowledge of when a node damages. We have modeled this finite sensitivity by applying non-zero false-negative attribution errors to our raw model FI [13]. This has no effect on the dynamics or on mortality, but does affect the FI. For any raw FI *f*_0_ = Σ_*i*_ *d*_*i*_/*n*; from *n* nodes, there are *n*_0_ = *f*_0_*n* damaged nodes. With a false-negative rate of *q*, *n*_*q*_ of these are overturned, where *n*_*q*_ is individually-sampled from a binomial distribution 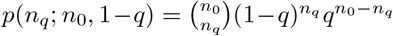. We use *f* = *n*_*q*_/*n* as the corrected individual FI. Since our model *f*_0_ tends to reach the arithmetic maximum of 1 at old ages, this effectively gives a maximum observed FI of 〈*f*_max_〉 = 1 − *q* [13]. We use *q* = 0.4 throughout.

Rather than using death ages, observational health data typically uses a binary mortality condition e.g. *M* = 0 if an individual is alive within 5 years of follow-up, or *M* = 1 otherwise. For ease of comparison between model and observational data, we adapt this approach in our analysis of mutual information [38, 39] between mortality and model nodes. This also lets us avoid the problems of individuals outlasting the length of an observational study and thus having no death age (which then requires dealing with “censored” data). Our entropy calculations will use binary entropy, *S*(*M*|*t*) = − *p*(0|*t*)log *p*(0|*t*) − *p*(1|*t*) log *p*(1|*t*), which we use to calculate information *I*(*M*; *D*_*i*_|*t*) = *S*(*M*/*t*) − *S*(*M*|*D*_*i*_,*t*). A similar type of analysis with information has been suggested by Blokh and Stambler [40].

We also compare our information theory results to a more standard survival analysis with hazard ratios [41]. The hazard ratio is the ratio of event rates for two values of an explanatory variable — e.g. with/without a deficit. A larger hazard ratio means a lower likelihood of surviving with the deficit than without. Hazard ratios are “semi-parametric”, since they extract the effects of variables on mortality rate from a phenomenological mortality model. We use the Cox proportional hazards model [42], which assumes exponential dependence of mortality rates. We show below that these survival analysis techniques are consistent with our non-parametric mutual information measures.

## III. RESULTS

We will focus on measures that can be compared between model and observational data, or that provide insight into the network structure of organismal aging. We start with model results, where we exactly know the network structure. We consider the age-structure in the damage, and then consider information measures. Then we explore how our results inform observational data. Finally we return to our model to test our insights by systematically manipulating the model network.

### A. Age-structure of damage

We construct two distinct FIs to capture the difference between well-connected hub nodes and poorly connected peripheral nodes. We measure low-degree damage by constructing *F*_low_ = Σ_*i*_ *d*_*i*_/*n* from a random selection of *n* = 32 nodes all with *k* = *k*_min_ = 2. Similarly, we measure high-degree damage with *F*_high_ from the top 32 most connected nodes (excluding the two most connected nodes, which are the mortality nodes).

The inset of Fig. 1 shows the average FI vs age for *F*_low_ and *F*_high_. We see 〈*F*_ow_〉 initially larger than 〈*F*_high_〉. Eventually with age, 〈*F*_high_〉 increases to match 〈*F*_low_〉 and even slightly exceed at very old ages. To better understand this age-structure we consider the connectivity of the damaged nodes.

Fig. 1 shows the cumulative average degree of damaged nodes 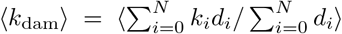 vs age *t*. Error bars represent the standard deviation between 10 different randomly generated networks. They are each comparable to or smaller than the point size, indicating that the age-structure represents the network topology rather than a single network realization.

For a uniform network or for damage rates independent of the degree of a node, we would expect 〈*k*_dam_〉 = 〈*k*〉 for all ages *t*. However, we see the average degree of damaged deficits start at 〈*k*〉, with an initial decrease until around age 25 and then an increase back to 〈*k*〉 — implying damage does not uniformly propagate through the network.

Initially damage is purely random, so 〈*k*_dam_(0)〉 = 〈*k*〉. Nodes with degree *k*_*i*_ < 〈*k*〉 are being damaged when 〈*k*_dam_〉/〈*k*〉 decreases from 1, and nodes of degree *k*_*i*_ > 〈*k*〉 are being damaged when 〈*k*_dam_〉/〈*k*〉 increases towards 1. 〈*F*_low_〉 is initially larger than 〈*F*_high_〉 in the inset of Fig. 1 because of the large amount of low-*k* damage at early ages.

We have developed a mean-field calculation in appendix A to understand the mechanism causing these results. To get the flavor of it, we can average the damage rates of a node over the possible states of its *k* neighbors- given their average damage *f*. From Γ_+_, we obtain the average damage rate

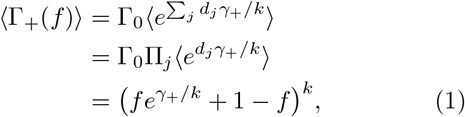

where the sum and product is over the *k* neighbors that are taken as independent — so that each neighbor node contributes the same factor to the result.

As shown in Fig. 2, these mean-field damage rates increase with smaller *k* at a given *f*. This results from Jensen’s inequality, since the damage rate is convex in the local frailty *f* and the lower degree nodes will have a broader distribution of local frailty for the same global frailty. This implies that low-*k* nodes should damage more frequently (until they are exhausted and *F*_low_ saturates). Indeed, in Fig. 3 the age-structure from the mean-field calculation shows the same early damage of low-*k* nodes shown in Fig. 1 and (in inset) the more-rapid growth of *F*_low_ compared to *F*_high_ at earlier times.

**FIG. 2.**
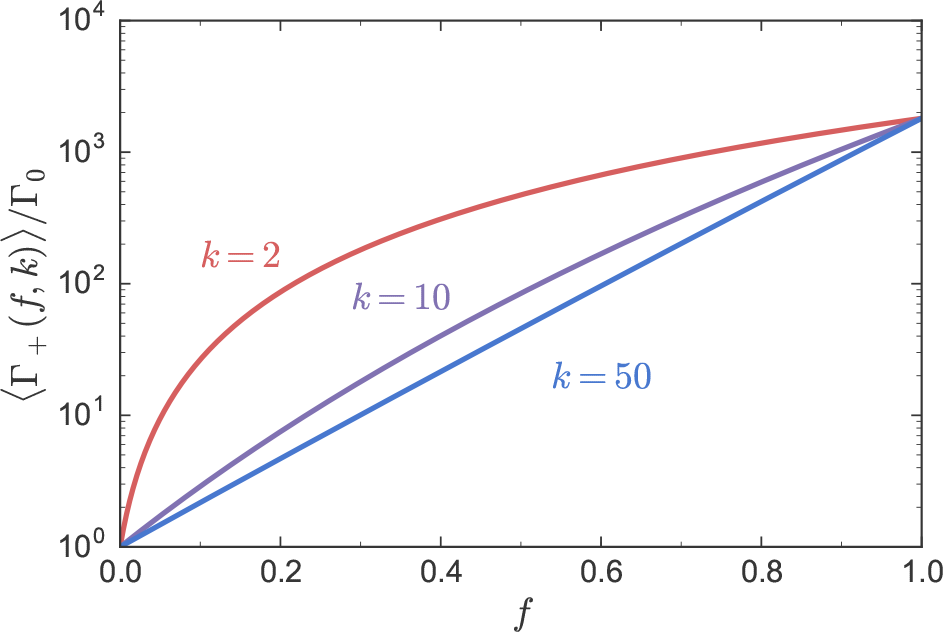
Average mean-field damage rates (Γ_+_)/Γ_0_ for nodes of a given degree *k* (as indicated) vs the local frailty of these nodes *f*, as given by Eq. 1. Low-connectivity nodes exhibit significantly higher damage rates at intermediate values of *f*.

**FIG. 3.**
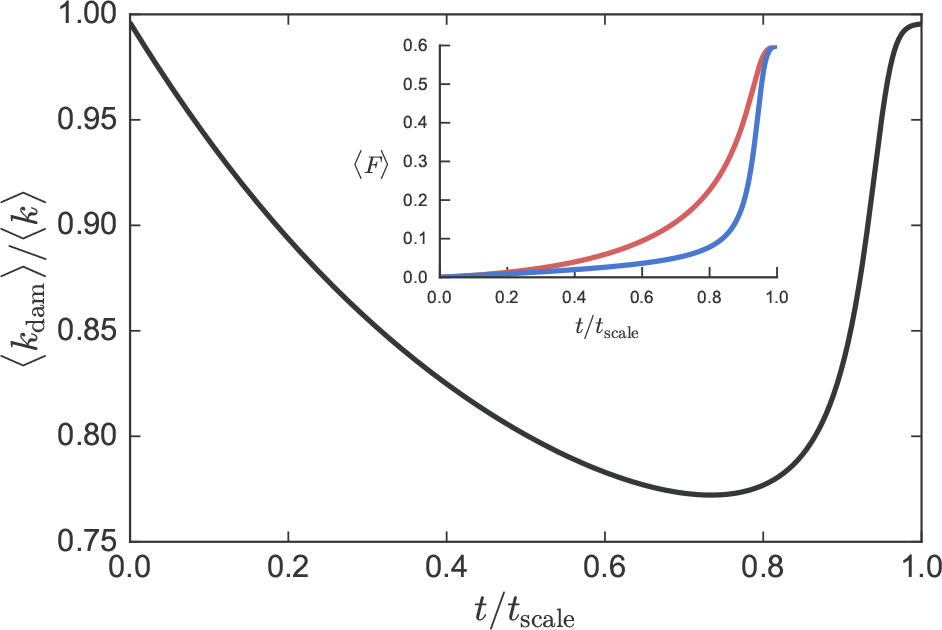
From our mean-field calculation in Appendix A, we show the average degree of damaged deficits 〈*k*_dam_〉 scaled by the average network degree 〈k〉 vs time scaled by the time when the network becomes fully damaged, *t*/*t*_scale_. The inset shows the average damage of high connectivity nodes 〈*F*_high_〉 in blue and low connectivity nodes 〈*F*_low_〉 vs the scaled time.

Interestingly, our mean-field calculation also shows a similar more-rapid growth of *F*_high_ compared to *F*_low_ at later times, as shown in the inset of Fig. 3. We believe that this largely is explained by the saturation of *F*_low_. However, as we explore below, there is also some heterogeneity among the nodes that are in *F*_low_.

### B. Node information with respect to mortality

Fig. 4 shows the mutual information between death age and individual nodes *I*(*A*; *D*_*i*_) for our model. Red points are a random selection of 100 low-connectivity nodes with *k* = *k*_min_ = 2, the blue points are the top 100 most connected nodes (excluding the 2 mortality nodes). For each selection, we have rank-ordered the nodes in terms of mutual-information. The mutual-information for both high and low connectivity nodes are comparable. This is surprising since previous work showed a monotonic increase of the average information with connectivity [13] — however that work used a different network for each individual, so that network properties other than the average connectivity were lost by pooling nodes of the same degree.

**FIG. 4.**
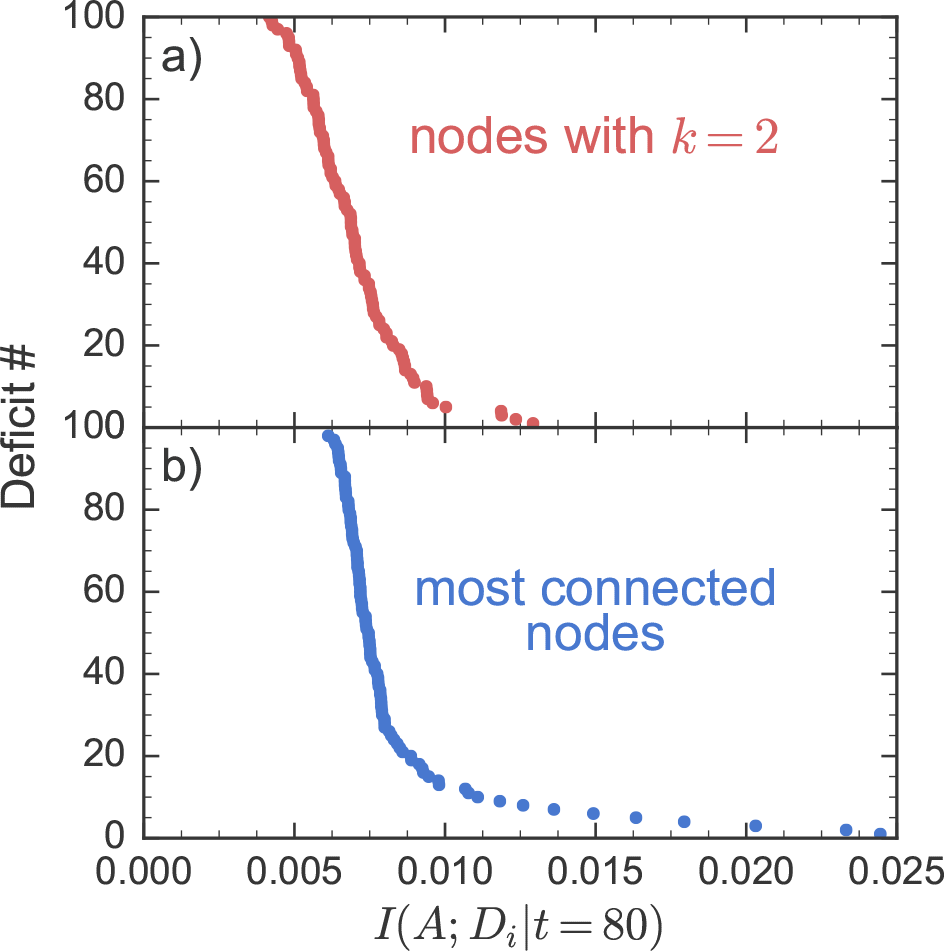
Mutual-information of selected model deficits *I*(*A*; *D*_*i*_\*t* = 80) at age 80 years, averaged over 10 randomly generated networks with 10^7^ individuals each and rank-ordered. Red points are low-*k* deficits, blue points are high-*k* deficits.

In Fig. 5, we show the “spectrum” of mutual information between death age and individual nodes *I*(*A*; *D*_*i*_|*t* = 80). We have used individuals at age *t* = 80 years, where the mutual information is close to maximal [13]. We use the same network for every individual, so that we do not lose the properties of the network between individuals. For the most connected nodes, in blue, we plot mutual information vs. the connectivity (degree *k*_*i*_) of the nodes. Here we see the monotonic trend of mutual information vs connectivity, though there is significant variation for individual nodes. For the least connected nodes, in red, all of the nodes have *k* = 2. Instead of connectivity, we considered the nearest neighbor degree 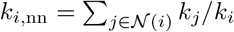 — i.e. the connectivity of the neighbors of a node. With respect to k_nn_, we see a similar monotonic increase of the mutual information for *k* = 2 nodes.

**FIG. 5.**
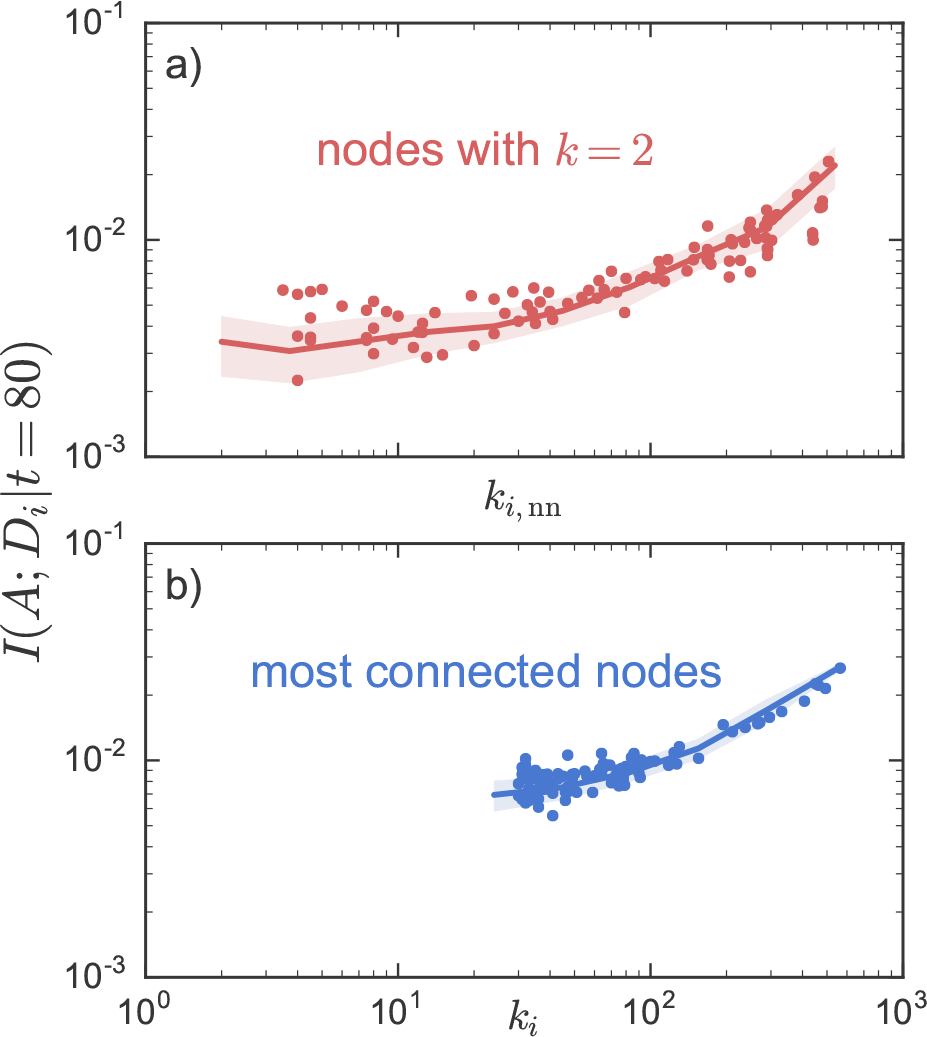
Model information spectra *I*(*A*; *D*_*i*_\*t* = 80) vs degree *k*_*i*_ for the top 100 most connected nodes in blue, or vs *k*_*i*,nn_ for a random selection of 100 peripheral nodes all with *k* = *k*_min_ = 2. Points show a sample of a single network, line shows an average over 10 randomly generated networks and the random choice of 100 nodes with *k* = 2, the shaded error region shows the standard deviation over the random networks.

Neighbor-connectivity *k*_nn_ is predictive of mortality for minimally connected nodes. We hypothesize that this is because the neighbor-connectivity affects when peripheral (*k* = 2) nodes are damaged, i.e. that peripheral nodes with low *k*_nn_ are damaged earlier than those with large *k*_nn_.

In the inset of Fig. 6 we confirm that high *k*_nn_ *k* = 2 nodes damage later. This allows high *k*_nn_ nodes to be informative of mortality because they are diagnostic of a more highly damaged network. From Fig. 6 we see that there is a large range of times for which lower-*k* nodes damage. Nevertheless, on average the high *k*_nn_ nodes at *k* = 2 damage before high-k nodes even though (see Fig. 5) they can be similarly informative.

**FIG. 6.**
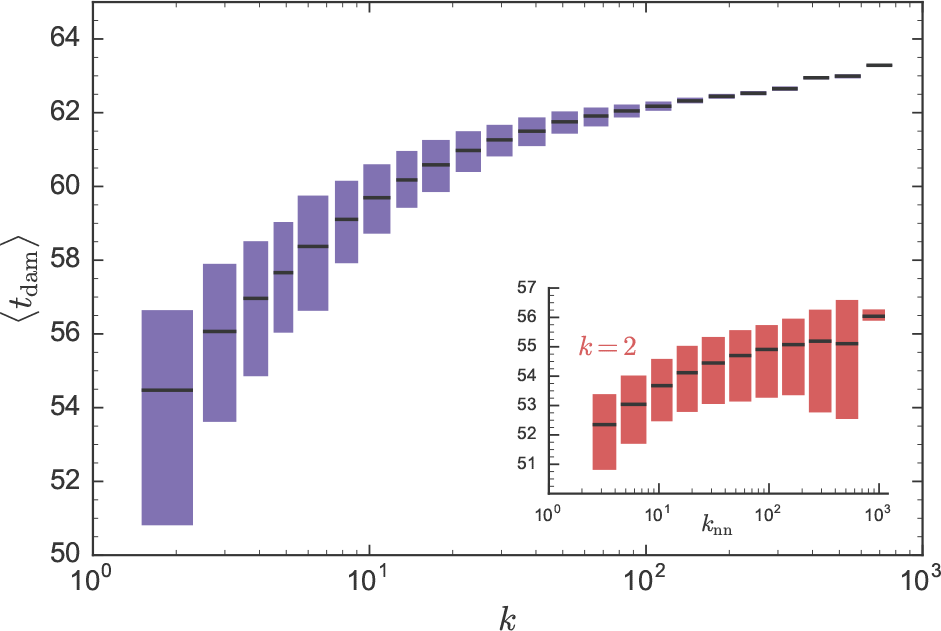
Average time of damage 〈*t*_dam_〉 vs degree *k* for all non-mortality nodes in the network. Inset shows 〈*t*_dam_〉 for *k* = 2 nodes vs nn-degree *k*_nn_. Nodes are binned based on *k*. The solid colored bars represent the entire range of average damage times observed for individual nodes within a bin, while the horizontal black lines indicate the average over the bin. All results are averaged for 10 randomly generated networks.

### C. Observational Data

Dauntingly, we have four challenges for assessing network properties from observational data: human studies are much smaller (typically with ≲ 10^4^ individuals) than our model populations so that results will be noisier, different studies will have quantitative differences due to cohort differences and choices of measured health attributes, we have no robust way of reconstructing networks from observed deficits so that the absolute connectivity of health-attributes is unknown (though attempts have been made [43]), and we have no map between specific observational attributes and specific model nodes.

However, we have observed qualitatively different model behavior from collections of highly connected vs peripherally connected nodes, and from peripheral nodes that are themselves connected to highly connected nodes vs those that are not. We expect such qualitative observations to be more robust to noise and cohort differences, and to the identification of individual attributes. Both laboratory test data and clinical assessments are available in large national studies of human aging. We expect that health-attributes assessed by laboratory tests are less connected than the high level functional attributes assessed clinically. We hypothesize that the corresponding FI measures (*F*_lab_ and *F*_clin_, respectively) should behave *qualitatively* like our model *F*_low_ and *F*_high_ — if the network underlying human aging has a similar structure as our model network.

From the American National Health and Nutrition Examination Survey (NHANES, see [44]), the 2003-2004 and 2005-2006 cohorts were combined, with up to 5 years of follow-up. Laboratory data were available for 9052 individuals and clinical data on 10004, aged 20+ years. Thresholds used to binarize lab deficits are found in [25]. From the Canadian Study of Health and Aging (CSHA, see [45]), 5 year follow-ups are obtained from 1996/1997. Laboratory data were available for 1013 individuals and clinical data for 8547, aged 65+ years. Thresholds used to binarize lab deficits are found in [21]. By approaching both the NHANES and CSHA studies with the same approaches, we can identify qualitatively robust features of both.

Fig. 7 shows the average FI vs age for *F*_lab_ in red and *F*_clin_ in blue for the NHANES in the main plot and CSHA in the inset. In both studies lab deficits accumulate earlier than clinical deficits. A crossover appears in the NHANES data around age 55 after which clinical deficits are more damaged than lab deficits. A similar crossover does not appear to happen in the CSHA data.

**FIG. 7.**
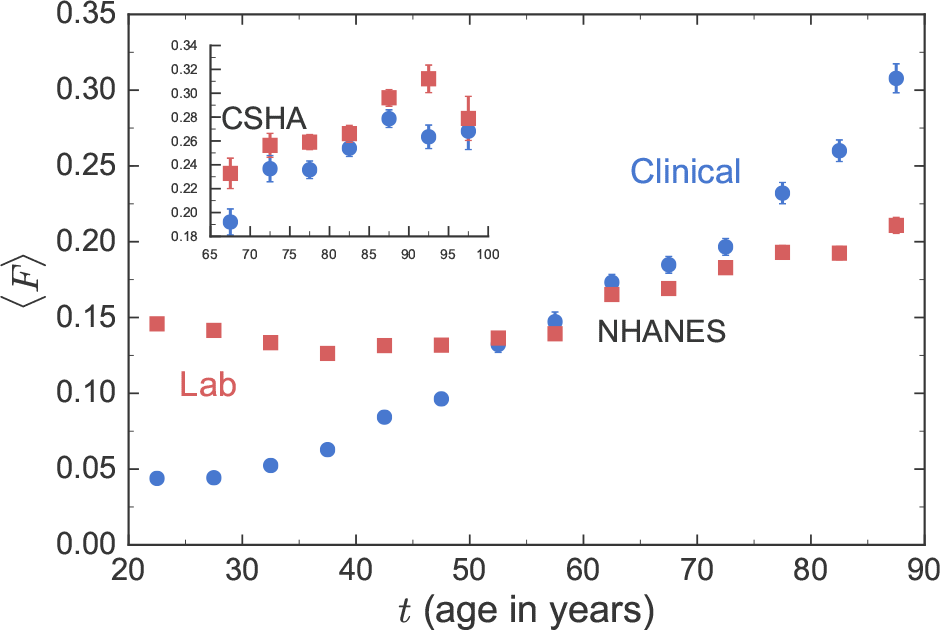
Average FI vs age *t* with 〈*F*_iab_〉 (red squares) and 〈*F*_clin_〉 (blue circles) from the NHANES dataset (main figure). The inset shows the same plot for the CSHA dataset. Error bars show the standard error of the mean. All individuals used in this plot have both *F*_clin_ and *F*_lab_ measured.

Comparable model results fors 〈*F*_low_(*t*)〉 and 〈*F*_high_(*t*)〉 were seen in Fig. 1. Thus, low-*k* nodes behave similarly to lab deficits, and high-*k* nodes behave similarly to clinical deficits in observational health data. Low-*k* nodes and lab measures both damage early and high *k* nodes and clinical measures both damage late.

Figs. 8 and 9 show deficits rank-ordered in information *I*(*M*; *D*_*i*_|*t*) for the NHANES and CSHA studies, respectively. These are information “fingerprints”. Red points correspond to lab deficits and blue to clinical deficits, as indicated. Both types of deficits have similar magnitudes of information, although clinical deficits are typically more informative. The comparable magnitudes of mutual information for the majority of individual deficits between lab and clinical FIs is consistent with earlier analysis that found similar association between lab and clinical FIs with mortality using survival analysis [21, 25, 27].

**FIG. 8.**
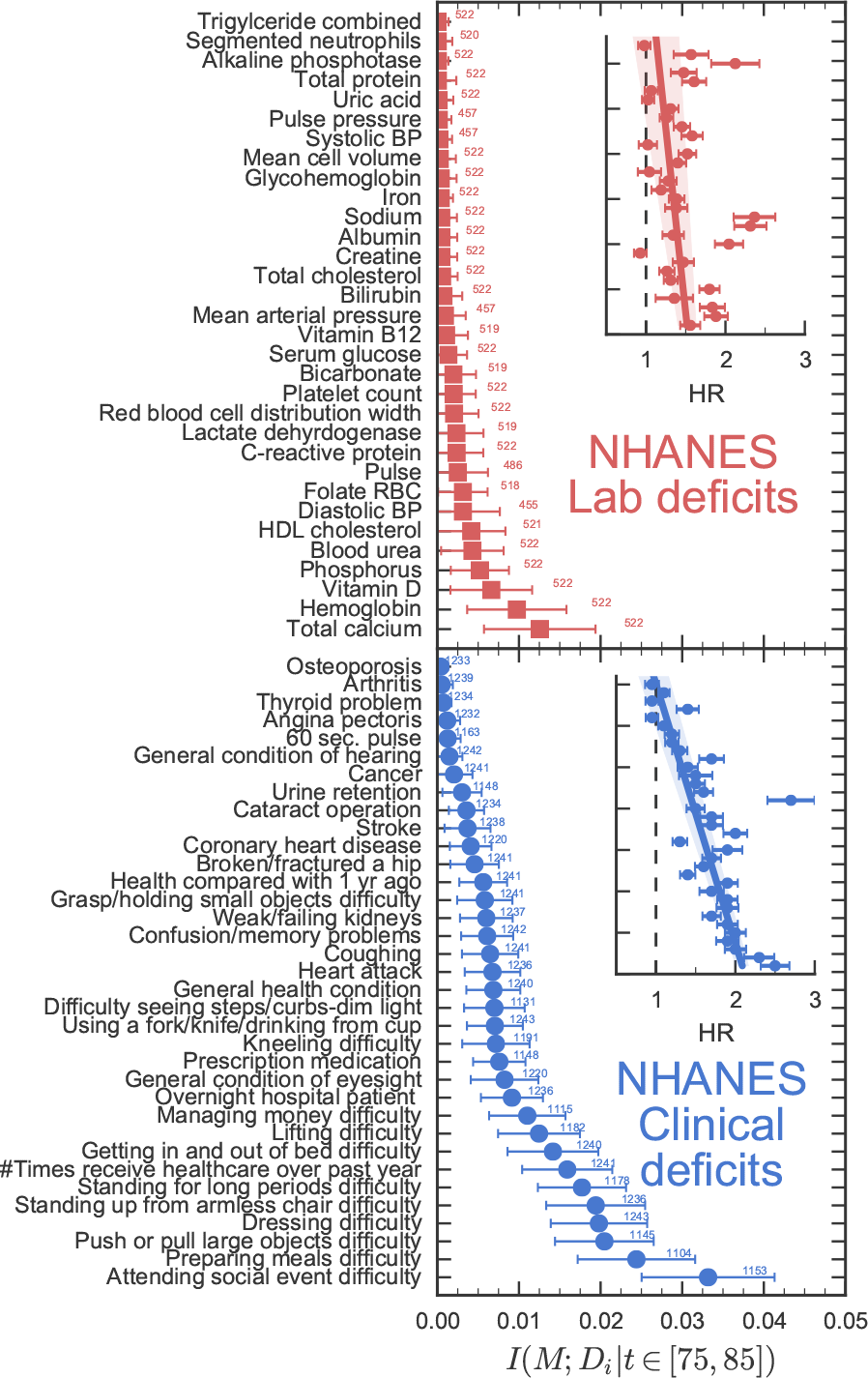
Rank-ordered deficits in terms of information *I*(*M*; *D*_*i*_\*t* ∈ [75,85]) for the NHANES dataset. Red points are lab deficits, blue points are clinical deficits. Error bars are standard errors found from bootstrap resampling. Small numbers next to the points indicate the number of individuals that were available in the data for the corresponding deficit. Insets show the corresponding hazard ratios for the deficits found from a Cox proportional hazards model regression, with the deficit and age used as covariates. The error bars show standard errors, and the line shows a linear regression through these points with the standard error in slope and intercept shown in a lighter color.

Insets in Figs. 8 and 9 show the corresponding hazard ratio (HR) for the deficit found from a Cox proportional hazards model regression, with the deficit value and age used as covariates. This traditional parametric analysis is often done with medical data [46]. The HR tends to increase as the rank-ordered information increases, indicating that our mutual-information approach is capturing similar effects. Nevertheless, we prefer mutual-information because it is non-parametric - i.e. it is not model-dependent - and so relies on fewer assumptions.

In Fig. 7, we showed that pre-clinical (lab) damage accumulates before clinical damage in observational data, and in Fig. 1 that low-degree damage similarly accumulates before high-degree damage in the model. Thus lab measures behave like model low-degree nodes and clinical measures behave like model high-degree nodes.

Our deficit-level analysis highlights the great variability of mutual information (and HR ratios) between individual deficits. We have shown that lab and clinical deficits have a range of mutual information, much like the model “spectra” shown in Fig. 5. We further note that the top 5 - 7 most informative clinical deficits in both the NHANES and CSHA datasets measure functional disabilities or dysfunction [47]. We find that these high level deficits are the most informative of mortality, and more informative than any of the lab deficits. From this, we hypothesize that highly informative clinical deficits will also be highly connected.

We have been able to partially reconstruct the network structure of clinical measures, as detailed and validated with our model in Appendix B. Our approach is based on the approach used to construct mutual information relevance networks [48, 49], and leads to reconstructed degrees 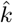 that are reliably rank-ordered for the highly connected nodes. (This approach did not reliably reconstruct properties of low-k nodes in our model, so we do not reconstruct networks for lab deficits.) In Fig. 10, we plot information with respect to mortality *I*(*M*; *D*_*i*_|*t* ∈ [75, 85]) for each deficit, where deficits are rank-ordered in terms of reconstructed degree 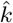. We see information increases with reconstructed degree for both the NHANES and CSHA clinical data. This shows that high information deficits correspond to high connectivity in both the observational and the model data. Also, nearly all of the functional disabilities hypothesized to have a high connectivity are also found to have a large reconstructed degree.

**FIG. 9.**
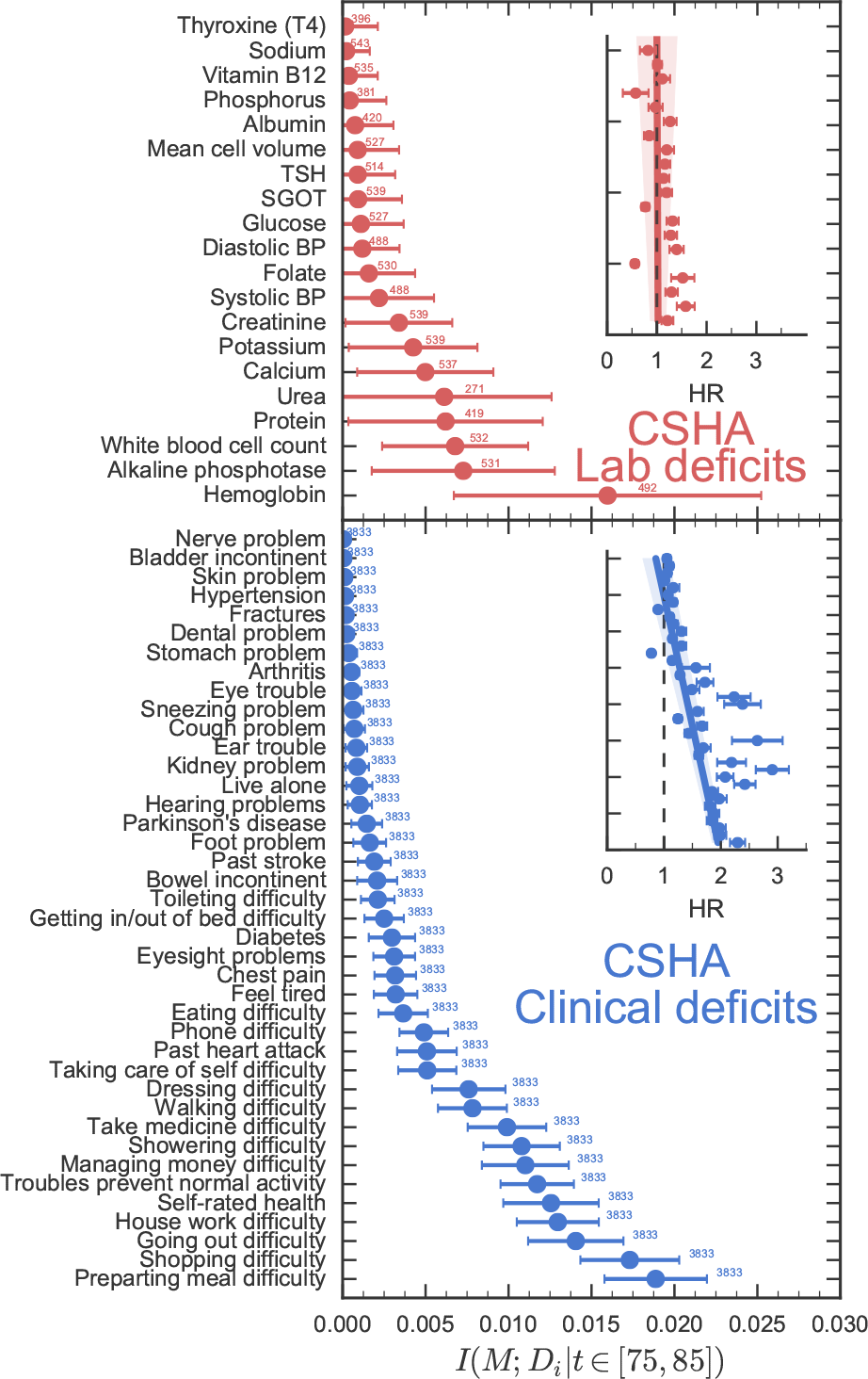
Rank-ordered deficits in terms of information *I*(*M*; *D*_*i*_\*t* ∈ [75, 85]) for the CSHA dataset. Red points are lab deficits, blue points are clinical deficits. Error bars are standard errors found from bootstrap re-sampling. Small numbers next to the points indicate the number of individuals that were available in the data for the corresponding deficit. Insets show the corresponding hazard ratios for the deficits found from a Cox proportional hazards model regression, with the deficit and age used as covariates. The error bars show standard errors, and the line shows a linear regression through these points with the standard error in slope and intercept shown in a lighter color.

**FIG. 10.**
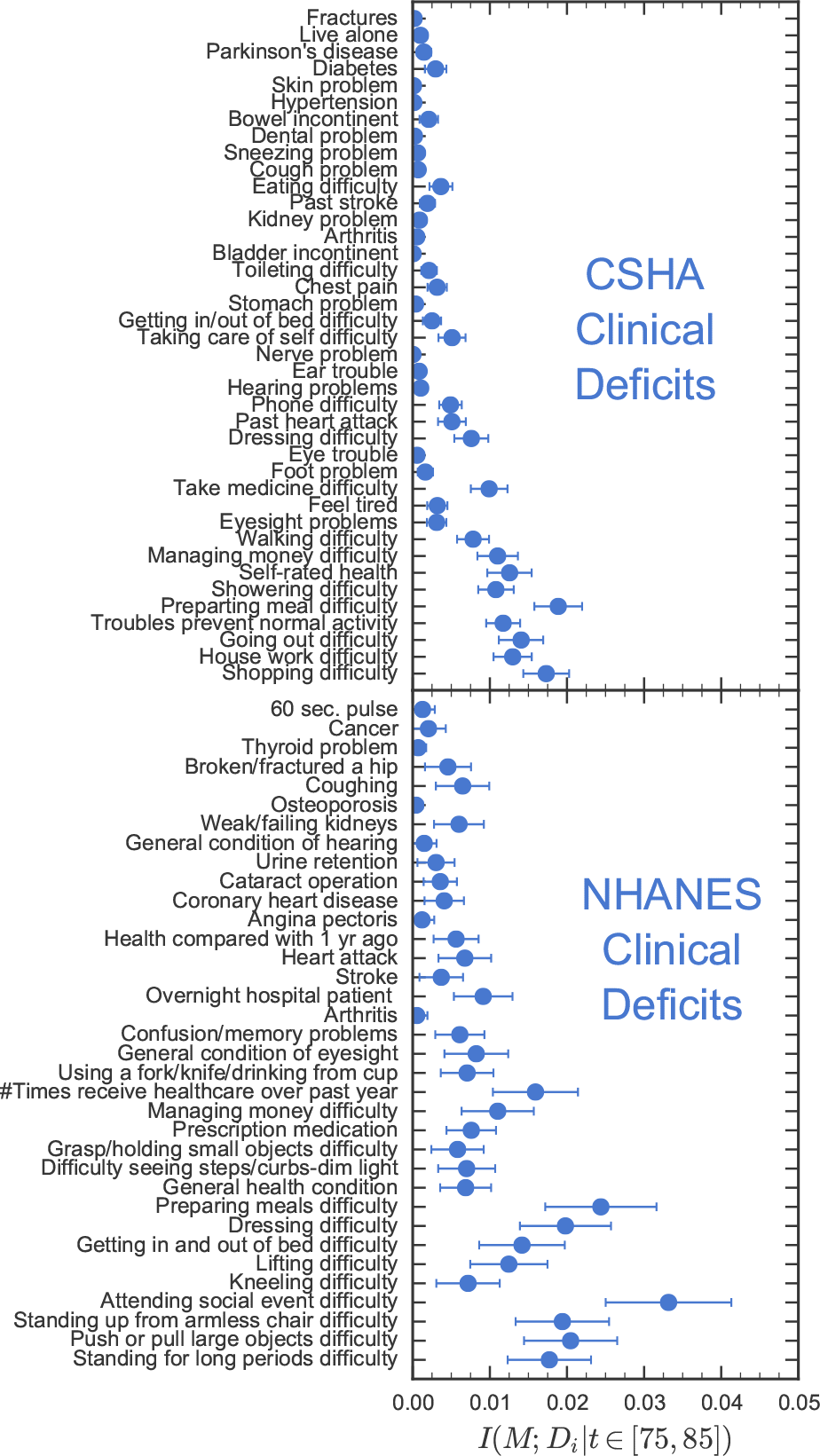
Rank-ordered clinical deficits in terms of reconstructed degree 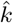 vs information with respect to mortality *I*(*M*; *D*_*i*_\*t* ∈ [75, 85]) for the NHANES and CSHA datasets. The reconstruction algorithm is detailed in Appendix B. Error bars are standard errors found from bootstrap re-sampling.

### D. Network structure

We have seen that our network model of aging is able to capture detailed behavior of lab and clinical FIs such as the the larger damage rates for low-*k* nodes at the same time as the surprising informativeness of some low-k nodes. The network is an important aspect of our model, and so far we have assumed that it is a preferential attachment scale-free network [29, 35, 36]. In this section, we show how specific properties of this network are necessary to obtain our results. We do this by exploring the qualitative behavior of different network topologies, and by identifying the network properties that alternative networks lack.

Our network model has predominantly disassortative correlations (due to the scale-free exponent α < 3 [50]) — meaning that low-*k* nodes tend to connect to high-*k* nodes, and that the average nn-degree decreases with degree [32]. We see this in Figure 11, where we plot the average nn-degree 〈*k*_nn_(*k*)〉 as a function of degree for our network. The purple points indicate our preferential attachment model network, and we see that the average nn-degree is inversely related to the degree. This shows disassortative correlations.

**FIG. 11.**
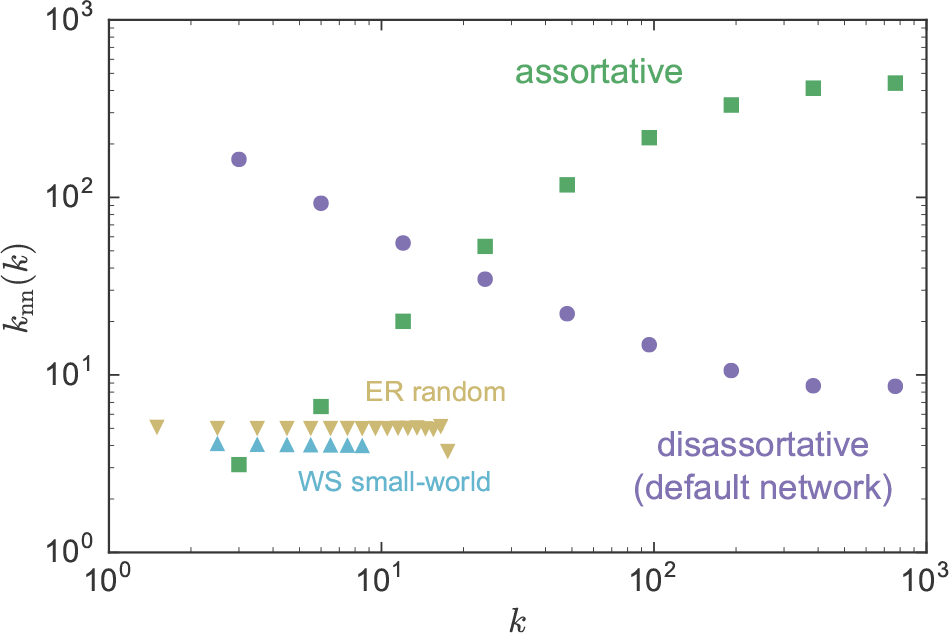
Average nn-degree 〈*k*_nn_(*k*)〉 vs degree *k* for a disassortative network (default network)(purple circles), an assortative network created by reshuffling the links (green squares) [51], a Erdős-Rènyi (ER) random network (yellow triangles), and a Watts-Strogatz (WS) small-world network (blue triangles). Note that 〈*k*_nn_(*k*)〉 is grouped into bins of powers of 2 and averaged within the bins for the scale-free networks. A bin for each degree is used for the ER random and WS small-world networks.

The green curve shows a rewired assortative network [32] made by preserving the degrees of the original network but swapping links. To do this we use the method of Brunet *et al,* using *N*^2^ rewiring iterations with a parameter *p* = 0.99 [51]. By modifying the nn-degrees of low degree nodes, we can investigate whether *k*_nn_ causes or is just correlated with informative low-*k* nodes. Note that we use only the largest connected component of the rewired network, with 〈*N*〉 = 9989 nodes over 10 network realizations.

The yellow triangles in Figure 11 show an ErdQs-Rènyi random network (ER). A random network is created by starting with *N* nodes, and randomly connecting each pair of nodes with probability *p*_attach_ = 〈*k*〉/(*N* − 1) [31]. This results in a (peaked) binomial degree distribution, and completely uncorrelated connections where *k*_nn_ = 〈*k*^2^〉/〈*k*〉 which is independent of individual node degree. As before, we only use the largest connected component, with 〈*N*〉 = 9805 nodes over 10 network realizations. The ER network also allows us to explore whether the heavy tail of the scale-free degree distribution is required to recover our observational results.

The light blue triangles in Figure 11 show a Watts-Strogatz (WS) small-world network [52]. This network starts with a uniform ring network with *k*_*i*_ = 〈*k*〉 for all nodes, and randomly rewires each link with probability *p*_rewire_ to another randomly selected node. We use *P*_rewire_ = 0.05 to get the effects of both high clustering (i.e. links between neighbors of nodes) and short average path-lengths between arbitrary pairs of nodes [31]. This network has a narrowly peaked degree distribution, with a rapidly decaying exponential tail. ER and WS networks are similar, as both have short average path lengths between arbitrary nodes and non-heavy-tailed degree distributions, but the WS small-world network also has high clustering for small *p*_rewire_.

To examine network effects on our network aging model, we have kept the same model parameters for the (default) preferential attachment disassortative network, the assortative network, the ER random network, and the WS small-world network. (The scale-free exponent *α* is only used in the disassortative and assortative networks.) We examine 10 random realizations of each network.

In Fig. 12 we show rank ordered information fingerprints for individual deficits *I*(*A*; *D*_*i*_\*t*, for the different network topologies as indicated. We observe striking differences in the scale and range of the mutual information with respect to mortality, and in the differences between the most and least connected nodes. The random and small-world network both have a significantly smaller scale of mutual information, together with a smaller range of variation.

**FIG. 12.**
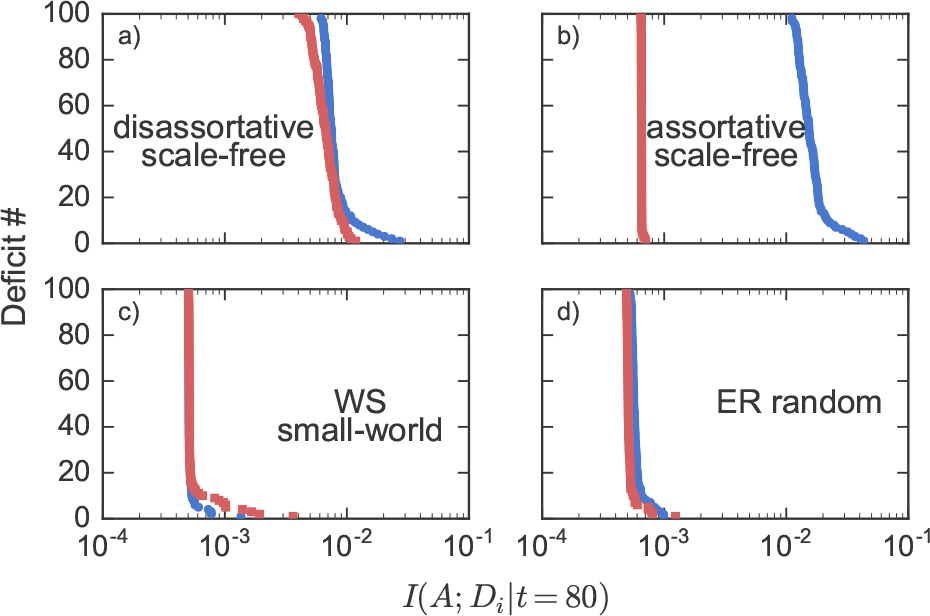
Rank ordered information *I*(*A*; *D*_*i*_\*t* = 80) for the different networks, as indicated. The top 100 most connected nodes in the network are in blue circles, and 100 randomly selected nodes of the lowest degrees are in red squares. Results for each different network topology are averaged over 10 randomly generated network realizations.

The scale-free disassortative (default) and assortative networks both have significantly higher scale of information for the most connected nodes, as well as considerable variation (approximately 10-fold) among them. However, while the disassortative network exhibits similar scales of information between the most and least connected nodes the assortative network does not. Furthermore, the assortative network shows only minimal variation of information among its least connected nodes.

Only the disassortative (default) network exhibits the fingerprint of mutual information exhibited by the NHANES and CSHA observational studies, in Figs. 8 and 9 respectively: with considerable variation of mutual information between deficits, overlapping ranges between lab (low) and clinical (high) connectivity deficits, and mutual information on the order of 10^−2^ for individual deficits.

We have also investigated the age-structure of the FIs generated by the low and high connectivity nodes for the different network topologies. In Fig. 13, we plot 〈*F*_low_(*t*)〉 vs 〈*F*_high_(*t*)〉 for the different network topologies. We see that the assortative network shows a rapid increase in *F*_low_, followed by growth of *F*_high_. In contrast, for the disassortative, random, and small-world networks there is comparable growth of both *F*_low_ and *F*_high_, though with higher *F*_low_ and a later cross-over for the disassortative network.

**FIG. 13.**
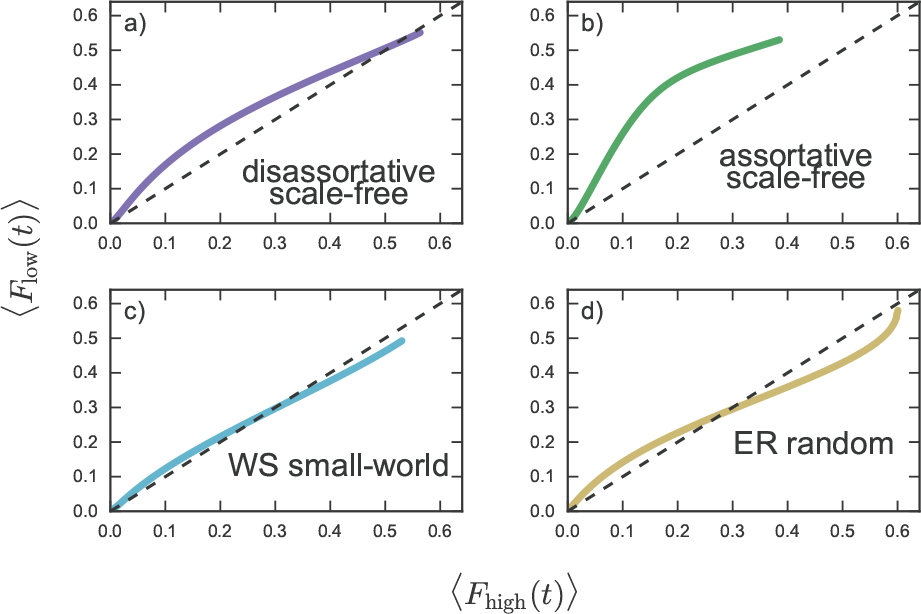
Average low-*k* 〈*F*_low_(*t*)〉 vs average high-*k* 〈*F*_high_(*t*)〉 plotted for *t* = 0 to *t* = 110 for our default network parameters (purple), the shuffled assortative network (green), the Erdős-Rènyi random network (yellow), and the Watts-Strogatz small world network (light blue). The dashed black line shows the line 〈*F*_low_(*t*)〉 = 〈*F*_high_(*t*)〉. Results are averaged over 10 randomly generated networks and the standard deviations are smaller than the line width.

From Fig. 12, we see that the early damage of *F*_low_ in the assortative network is associated with significantly smaller mutual information of low-*k* nodes. Our dynamical mean-field theory (MFT) in Appendix A allows us to narrow down what aspects of the network are leading to this behavior. In the MFT the only aspects of the network included are the degree distribution *P*(*k*) and nn-degree correlations *P*(*k*|*k′*).

From our MFT, in Fig. 14 we show the average low-*k* FI vs the average high-*k* FI, 〈*F*_low_(*t*)〉 vs 〈*F*_high_(*t*)〉. In purple we use the (default) preferential attachment disassortative correlations, in green we use assortative correlations (Eq. A8), and in light blue we use a WS small-world network. We see qualitative agreement with the age-structure shown in Fig. 13 - confirming that nn-degree correlations (included in our MFT) are important for the observed age-structure.

**FIG. 14.**
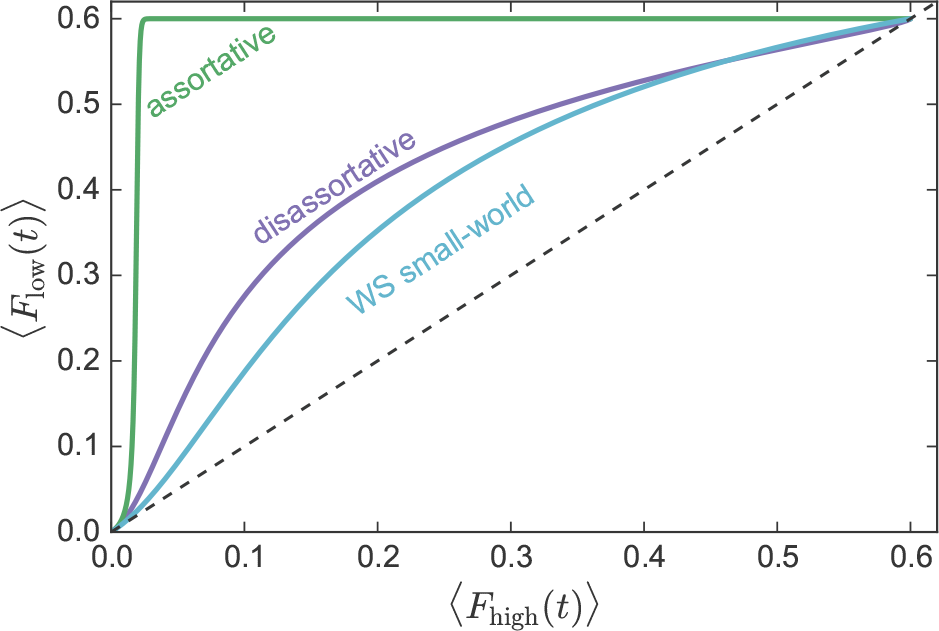
Average low-*k* 〈*F*_low_(*t*)〉 vs average high-*k* 〈*F*_high_(*t*)〉 from our mean-field model in Appendix A. The dashed black line shows the line 〈*F*_low_(*t*)〉 = 〈*F*_high_(*t*)〉. A scale-free network with preferential attachment disassortative correlations (default network) in purple, scale-free network with assortative correlations in green, and a WS small-world network with neutral correlations in light blue.

## IV. DISCUSSION

In our model low-*k* nodes tend to damage before high-*k* nodes. This is because of the larger average damage rates of low-*k* nodes compared to high-*k* nodes (Fig. 2). At the same time, our information spectrum shows that information *I*(*A*; *D*_*i*_\*t* increases with *k*. Roughly speaking, high-*k* nodes need a larger local frailty *f* to have comparable damage rates as low-*k* nodes. Thus, damage of high-*k* nodes is informative of high network damage, which also leads to mortality. This is why high-*k* nodes both damage later and are informative of mortality (Fig. 5b).

However, some low-*k* nodes also damage later and are highly informative of mortality. Information *I*(*A*; *D*_*i*_|*t*) increases with *k*_nn_ for the low-*k* nodes, and low-*k* high-*k*_nn_ nodes damage later. This can be explained using the network structure. low-*k* nodes are protected from damage when they are connected to high-k nodes. Rapidly damaging low-*k* nodes without this protection tend to damage early for most individuals, giving these nodes a low information value of mortality. Conversely, protected nodes tend to damage only when their high degree neighbors start to damage, which only occurs when the network is heavily damaged and close to mortality. As a result, only the low-*k* nodes with high-*k*_nn_ are highly informative (Fig. 5a). Interestingly these nodes still tend to damage before high k nodes, leading to an early predictor of mortality.

Degree correlations control the average degree of neighboring nodes and hence control the amount of protection in low-*k* nodes. By modifying the degree correlations in the network in our computational model we have shown that this protection can be caused by disassortative correlations — where low-*k* nodes tend to attach to high-*k* nodes. Conversely, eliminating low-*k* high-*k*_nn_ nodes by modifying the network to introduce assortative correlations removes this protection, and we then find all low-*k* nodes have low information (Fig. 12b).

Our mean-field model allows us to explicitly modify the degree distribution and the degree correlations with the nearest-neighbor degree distribution *P*(*k′*|*k*), and to include no other network features. In our mean-field model we see similar results to our computational model where, e.g., adding assortative correlations increases the rate at which *F*_low_ increases with respect to *F*_high_. This confirms that degree distribution and degree correlations largely determine the early damage of low-*k* nodes that we observe in scale-free networks.

Degree distributions and correlations only weakly control the behavior of ER random and WS-small world networks. The low variation in *k* and *k*_nn_ in those networks results in a lack of contrast between the damage rates of nodes. This leads to node information that is nearly constant throughout the network and to only small differences in the damage structure of low-*k* and high-*k* nodes (Fig. 12c and d). This also leads to low magnitude of the mutual information per node, since nodes behave much more uniformly and “randomly” than in a scale-free network. However, we can still see some protection in low-*k* nodes. This is particularly apparent in the ER random network when *F*_high_ surpasses *F*_low_ (Fig. 13d).

The observational FIs, *F*_clin_ and *F*_lab_, measure distinct types of damage — clinically observable damage that tends to occur late in life and pre-clinical damage that is typically observable in lab tests or biomarkers before clinical damage is seen. These different types of damage are useful, because they occur at different times during the aging process though both are similarly informative of mortality. We have shown that two different FIs, *F*_clin_ and *F*_lab_, are analogous to the model *F*_high_ and *F*_low_ respectively. In our model, low-*k* deficits damage first and lead to the damage of high-*k* nodes. In addition, both are similarly informative of mortality although with some high-*k* deficits somewhat more informative.

Furthermore, the behavior of observational deficits seems to best resemble the behavior of the computational model with a scale-free network and disassortative correlations. Node information seen in the (default) scale-free disassortative network is a much better qualitative match of observational data, as compared with scale-free assortative, WS small-world, or ER random networks. The age-structure of damage also qualitatively agrees with this topology.

Our analogy between observational deficits and model nodes allows us to make predictions about the underlying network structure of observational health deficits, even though we cannot directly measure this network. The observational network should have a heavy-tail degree distribution, so that a large range of possible information values can be obtained. The network should also include disassortative correlations so that there are connections between high-*k* and low-*k* nodes, allowing low-*k* nodes to be informative of mortality.

Indeed, from observational data we find that clinical deficits that integrate many systems into their performance (e.g. functional disabilities, or social engagement) are very informative (Figs. 8 and 9). In contrast, single diagnoses, even ones strongly associated with age such as osteoporosis, on their own offer less value. The model interpretation of this is that these high information disability deficits have a higher connectivity than lower information clinical deficits. It intuitively makes sense for deficits that integrate many systems to have a large connectivity. In support of this, our partial network reconstruction (Fig. 10) shows that high information clinical deficits in both the NHANES and CSHA correspond to nodes with a high reconstructed degree.

We have kept the model parameterization unchanged from the default parameters. This has allowed us to explore the impact of network topology on mortality statistics (a small effect) and on mutual information between health deficits (a qualitatively strong and distinctive effect). The *F*_high_ and *F*_low_ model phenomenology are qualitatively affected by changes in network topology. This indicates that both *F*_high_ and *F*_low_ are usefully distinct characteristics of health in our network model. We believe that *F*_clin_ and *F*_lab_ are similarly useful and distinct characteristics of human health. This has been a focus of recent studies [21, 22, 27], and we believe our results give insight into the mechanisms of these different health measures.

## A. Acknowledgments

We thank ACENET and Compute Canada for computational resources. ADR thanks the Natural Sciences and Engineering Research Council (NSERC) for operating grant RGPIN-2014-06245. KR is funded in this work by career support as the Kathryn Allen Weldon Professor of Alzheimer Research from the Dalhousie Medical Research Foundation, and with operating funds from the Canadian Institutes of Health Research (M0P-102544) and the Fountain Innovation Fund of the Queen Elizabeth II Health Science Foundation. SGF thanks NSERC for a CGSM fellowship.

## APPENDIX A: Mean-field theory of network model

### a. MFT introduction

In this appendix we present details of our mean-field theory (MFT) analysis of our network model, which is based on earlier work on epidemic processes in complex networks [53] and uses ideas from Gleeson [54] to include mortality dynamics. Our MFT here is more complicated than a very simple MFT presented in our earlier work [13], but captures much more of the aging phenomenology. (The MFT here reduces to the simpler MFT when all of the nodes have the same degree.)

By MFT we mean a set of deterministic dynamical equations for damage probabilities of network nodes, including mortality nodes. These coupled ordinary differential equations (ODEs) are solved with standard numerical integrators, and the solutions used to generate, e.g., 〈*F*_low_(*t*)〉 and 〈*F*_high_(*t*)〉. Significantly, in our MFT we retain the full degree distribution *P*(*k*) and degree correlations *P*(*k*|*k′*) of our stochastic network model. This allows us to identify what model behavior is controlled by the degree distribution and degree correlations.

### b. MFT calculation

We average the damaged probabilities *p*(*d*_*i*_ = 1) and the undamaged probabilities *p*(*d*_*i*_ = 0), conditioned on the damage of the mortality nodes, over all nodes of the same degree *k*:

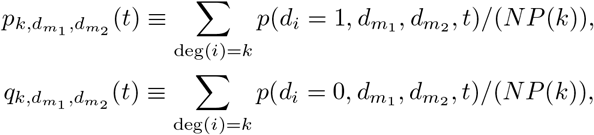

where the mortality states are indicated by 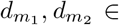 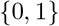, *N* is the number of nodes, and *P*(*k*) is the degree distribution. The resulting joint probabilities are 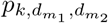 and 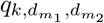, for damaged and undamaged nodes respectively. These joint probabilities satisfy

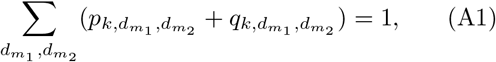

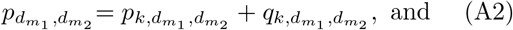

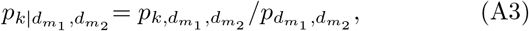

where the first equation is a normalization condition, the second completeness, and the third Bayes’ theorem for conditional probabilities. From our mortality rule of 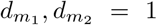, the probability of mortality is *p*_dead_ = *p*_*k*,1,1_ + *q*_*k*,1,1_, for any *k*.

The probability of a neighbor of a node of degree *k* being damaged (which is its local frailty *f*) given a particular mortality state is

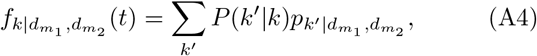

where *P*(*k′*/*k*) is the conditional degree distribution, or “nearest-neighbor” degree distribution. *P*(*k′*/*k*) describes the structure of connections in the network, and can be varied independently of the degree distribution *P*(*k*).

Writing exact master equations for *N* nodes is impractical since there would be 2^*N*^ distinct states to track, with even more distinct transition rates. As an enormous simplification, we use averaged damage and repair rates of nodes of a given connectivity *k*. This is our key mean-field simplification. To do this we approximate 〈*d*_*i*_*d*_*j*_〉 = 〈*d*_*i*_〉 〈*d*_*j*_〉 for all nodes, and approximate the number of damaged neighbors by a binomial distribution 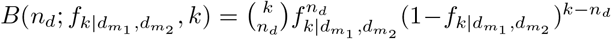 where the average proportion of damaged neighbors will be 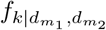. Using Eq. A4, we can then calculate our MFT damage and repair rates,

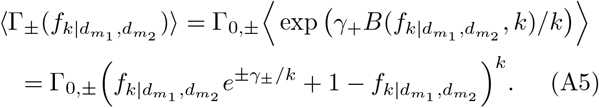

These rates correspond to Eq. 1 and are shown in Fig. 2.

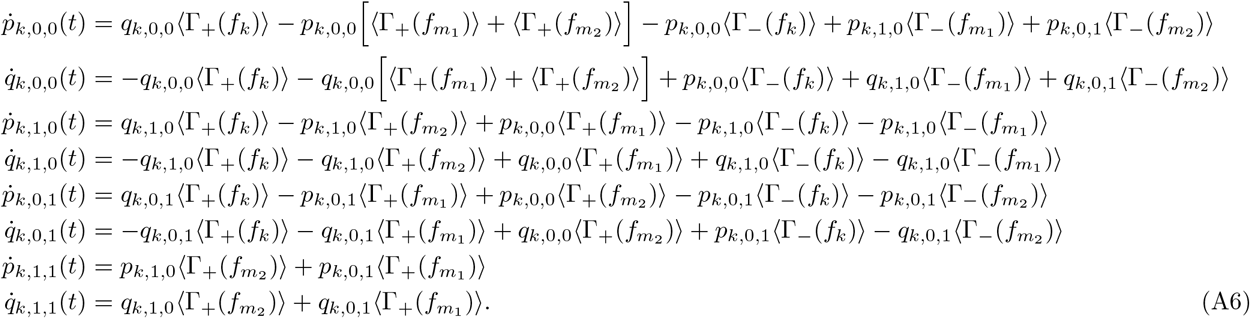

In these equations we have not shown the mortality state indices of *f*_*k*_ for readability, but they are the same as the associated *P* or *q* factors. We have also defined 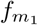 and 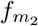 as the local frailties of the first and second mortality node, respectively. We have 8 equations for each distinct degree *k*. The last two equations determine the mortality rate, 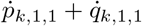.

The mean-field model couples the dynamics of the lowest degree (*k* = 2) with all degrees up to the two highest (mortality nodes). Solving the equations requires us to explicitly determine the two mortality node degrees. While approximate calculations of the maximum degree of scale-free networks are available [55], we need the two highest degrees. We use 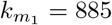 and 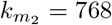, based on the averages from simulations of the network. Similarly, we use 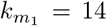 and 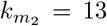 for ER random networks and 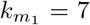 and 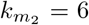 for WS small-world networks. Qualitatively, our qualitative MFT results do not depend on these mortality node degrees, as long as they are sufficiently large. The minimum degree *k*_min_ is determined by the network topology.

### c. MFT networks

The degree distribution *P*(*k*) and conditional degree distribution *P*(*k′*|*k*) represent the network structure in our MFT. The exact *P*(*k*) for our default shifted-linear preferential attachment networks [36], ER random networks, and WS small-world networks [56] are known. (We remove zero degree nodes from the ER random degree distribution, so that *P*_k≠0_(*k*) = *P*(*k*)/Σ_*l*≠0_ *P*(*l*).) Using various *P*(*k′*|*k*) we can then put different degree correlations into our MFT network. We include three While the node degree is explicit in Eq. A5, the degree correlation is still included through the average local damage in Eq. A4.

Using these average damage/repair rates as transition probabilities, we can write a master equation for nodes with connectivity 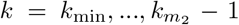 and given the global state of the mortality nodes: types of degree correlations, uncorrelated (neutral), assortative, and disassortative [31].

For a network with uncorrelated (neutral) connections,

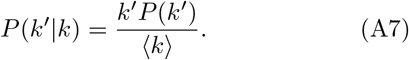

We then have 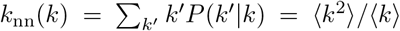, so that all nodes have the same nn-degree. These correlations are used for ER random and WS small-world networks. (Although these are not the exact correlations for a WS small-world network, they recover the approxi-
mately constant knn that we observed in Fig. 11.)

In a network with assortative correlations, nodes tend to be connected to other nodes of similar degree. Assortative correlations that approximate those used in our computational model in Sec. IIID are [57]

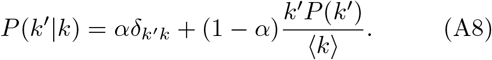

These lead to,

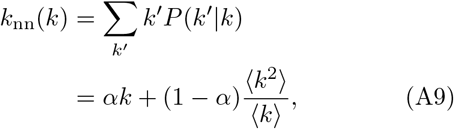

which increases linearly with *k* (see Fig. 11). Changing α modifies the amount of assortative correlation; we use α = 0.8.

In a network with disassortative connections, nodes tend to be connected to other nodes of differing degree. The (disassortative) correlations for our default shifted-32
where m = (k}/2 = k_min_ and A = m(a — 3). This is exact in the limit N → ∞TO [36],

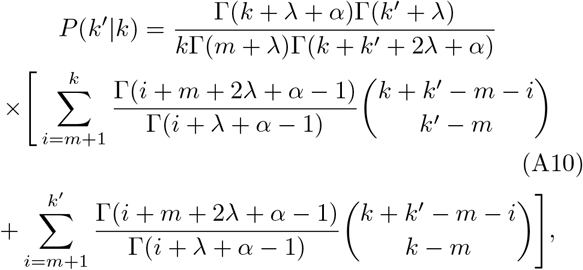

where *m* = 〈*k*〉2 = *k*_min_ and λ = *m*(α − 3). This is exact in the limit *N* → ∞ [36], and gives disassortative correlations where *k*_nn_(*k*) decreases with *k* (as illustrated in Fig. 11).

We can use our mean-field model for uncorrelated, assortative, and disassortative networks with any degree distribution *P*(*k*), using these various *P*(*k′*|*k*).

### d. MFT results

We numerically solve Eq. A6 for the probabilities 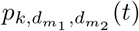 and 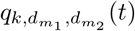. These then allow us to calculate the average FI,

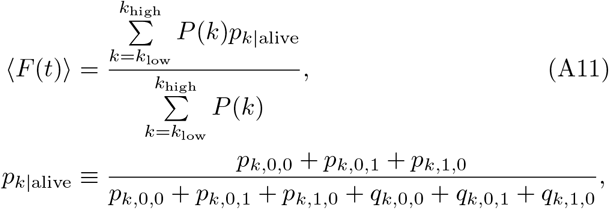

where the average is over the surviving individuals. The *k*_low_ and *k*_high_ determine the nodes included in the FI. For *F*_high_, we choose 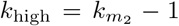 and *k*_low_ so that 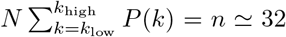 for the smallest possible *k*_low_ (32 is the number of FI nodes typically used in our model and observational studies). For *F*_low_, we choose *k*_low_ = *k*_min_ and choose the smallest *k*_high_ so that n ≃ 32.

We also calculate

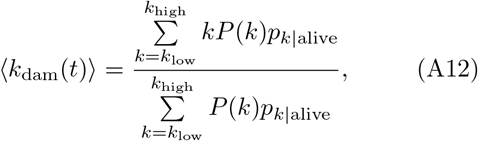

which is the cumulative average degree of damaged nodes. This is a measure of network damage.

In Fig. 15 we plot the average mortality rates vs age fro different network topologies, with colored circles showing the computational model results and colored lines for the corresponding mean-field model results. Black squares indicate observed mortality rates [58]. Most of the network topologies, with the exception of the assortative scale-free network, approximately captures the observational Gompertz’ law at later ages. This indicates that mortality data alone does not strongly constrain the network topology.

**FIG. 15.**
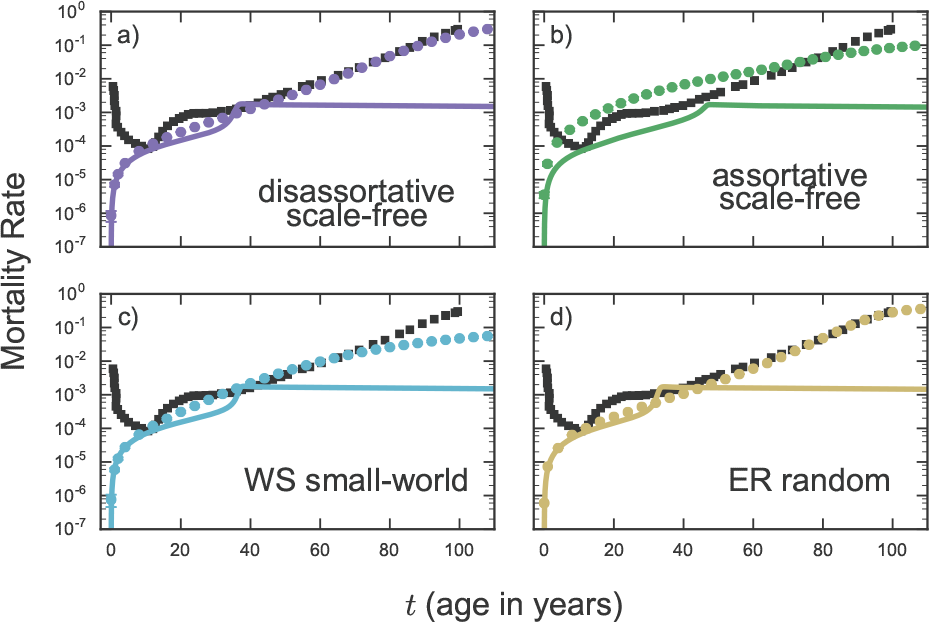
Mortality rate vs age for each of the networks. a) Disassortative scale-free network (purple circles), b) assortative scale-free network (green), c) WS small-world network (light blue), and d) ER random network (yellow). Circles show the computational model and lines show the mean-field model. Computational results are averaged over 10 randomly generated networks and error bars show the standard deviations. Black squares are observed human mortality rates [58].

The mean-field and computational models agree well for low *t*, albeit in a model regime that does not include developmental or child-mortality effects [13]. Interestingly, the mean-field models all exhibit an anomalous saturation of mortality rates that occurs around 30 - 50 years. This saturation arises from faster damage rates in the mean-field models. The mechanism is simple: rapidly damaging nodes drop out of the full model once they are damaged, but continue to contribute to the average damage rates in the mean-field model through Eq. A5. Accordingly, we plot most of our MFT results vs *t*/*t*_scale_ — where *t*_scale_ is the time that the network is fully damaged, with *F*_Scale_ = 1 − *q*.

Note that we have not shown our MFT results for the ER random network in Fig. 14. This is because our MFT behaves poorly when it includes nodes with k ≤ 2 (see Fig. 11), due to their great variability of local frailty *F*_i_.

## APPENDIX B: High-fc Network Reconstruction

To reconstruct network connections from observed states of nodes, we use the state of each deficit (node) at a given age *t* (or narrow range of ages in observational data) for each individual in the sample, and calculate the mutual information between individual deficits, *I*(*D*_*i*_; *D*_*j*_|*t*). Connections in the model create correlations between nodes, so a large *I*(*D*_*i*_; *D*_*j*_|*t*) could indicate a connection. We use data where individuals are the same age (or ± 5 years in observational data), so that time is not a confounding variable. Nevertheless, determining whether a given connection exists or not requires a threshold on *I*(*D*_*i*_; *D*_*j*_|*t*). We would only assign a connection between nodes if the mutual information is above this threshold. However, we have no practical way of determining such a threshold; a similar challenge exists when considering temporal data [59].

In preliminary tests with our model we have found that matching the reconstructed average degree with the exact average degree is a reliable way of determining a threshold (data not shown), but we still have no way of determining the average degree from observational data. Instead, we use a simple parameter-free method borrowed from mutual-information relevance networks [48, 49]. We calculate a “reconstructed” degree by adding the information for each possible connection to the node in the network, 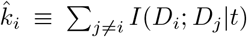 [60]. For nodes that aren’t connected, *I*(*D*_*i*_; *D*_*j*_|*t*) ≈ 0, while *I*(*D*_*i*_; *D*_*j*_|*t*) is large for connected nodes. While we cannot reconstruct the actual network, we can reconstruct the rank-order degree of high-*k* nodes as long as 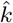 is roughly monotonic with the actual degree *k*.

In Fig. 16, we have validated this approach with the top 32 most-connected model nodes. We use 10000 individuals for our validation, approximately the same number of people we have available in the larger NHANES and CSHA observation studies that we consider. We know that our model information tends to increase with degree for the high degree nodes, see Fig. 5. Fig. 16 shows that information also increases with the reconstructed degree 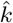. The inset showing *k* vs 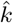 indeed shows that the reconstructed degree is approximately monotonic with the exact degree — especially at higher *k*.

**FIG. 16.**
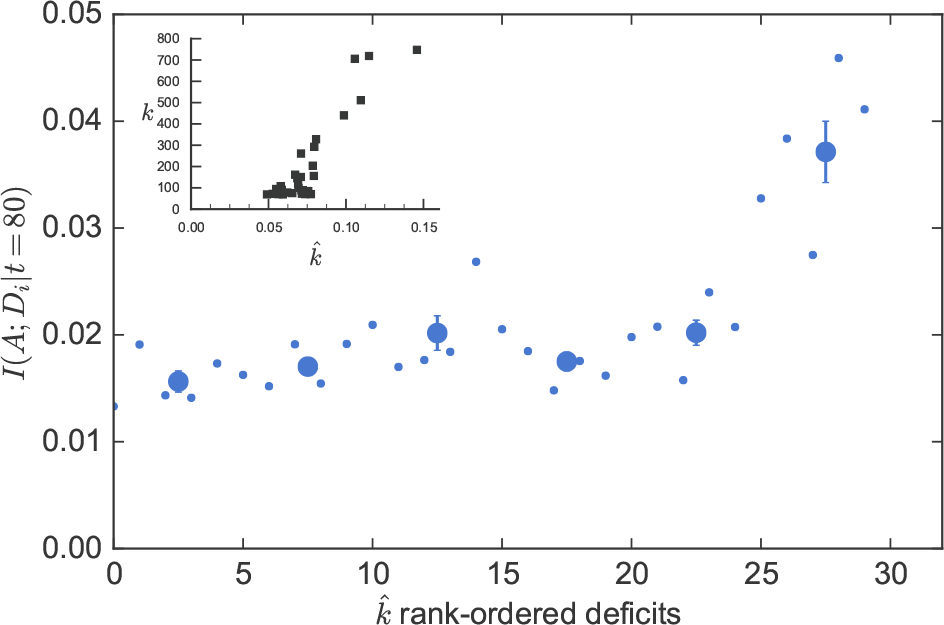
Information *I*(*A*; *D*_*i*_\ = 80) vs rank-ordered deficits using reconstructed degree 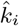, for our computational model. The top 32 most connected nodes are reconstructed with 10000 individuals. The smaller (blue circles) points show individual nodes, the larger points show a binned average, and error bars are the standard error of the mean within each bin. The inset (black squares) shows the exact degree *k* vs the reconstructed degree 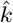.

This means the reconstructed degree should provide a reasonable rank-order degree for observational data, see e.g. Fig. 10. Nevertheless, low-degree nodes are not reliably rank-ordered. Accordingly we only reconstruct clinical 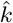 with this approach.

